# HIV lymphoid tissue fibrosis occurs in the earliest stages of acute HIV infection and is associated with macrophage-derived TGF-β

**DOI:** 10.64898/2026.02.02.703316

**Authors:** Jodi Anderson, Carlo P. Sacdalan, Somchai Sriplienchan, Nittaya Phanuphak, Kevin Escandón, Garritt Wieking, Erika Helgeson, Caitlin David, Jeffrey G. Chipman, Alexandra Schuetz, Sandhya Vasan, Daniel C. Douek, Timothy W. Schacker

**Affiliations:** Division of Infectious Diseases and International Medicine, Department of Medicine, University of Minnesota, Minneapolis, MN, USA; SEARCH Research Foundation, Bangkok, Thailand; Faculty of Medicine, Chulalongkorn University, Bangkok, Thailand; Institute of HIV Research and Innovation, Bangkok, Thailand; Division of Biostatistics and Health Data Science, University of Minnesota, Minneapolis, MN, USA; Department of Surgery, University of Minnesota, MN, USA; U.S. Military HIV Research Program, Center for Infectious Disease Research, Walter Reed Army Institute of Research, Silver Spring, MD, USA; Henry M. Jackson Foundation for the Advancement of Military Medicine, Inc., Bethesda, MD, USA; Armed Forces Research Institute of Medical Sciences, Bangkok, Thailand; Vaccine Research Center, National Institute of Allergy and Infectious Diseases, National Institutes of Health, Bethesda, MD, USA

**Author notes:** Address correspondence to: Timothy W. Schacker, Division of Infectious Diseases and International Medicine, Department of Medicine, University of Minnesota, Minneapolis, MN, USA.

**Keywords:** HIV, TGF-β, macrophage, lymphoid tissue fibrosis

## Abstract

HIV infection is associated with lymphatic tissue damage caused by collagen deposition in the fibroblastic reticular cell network (FRCn) of the parafollicular T cell zone (TZ), which leads to impaired antigen presentation, T cell depletion, and reduced antibody formation. In treated chronic HIV infection, damage persists despite antiretroviral therapy (ART), but it is not known if very early initiation of ART could limit damage and preserve immune function. We studied participants in the Thai RV254 acute infection cohort and found significant collagen deposition in the FRCn even in the earliest Fiebig stage 1 acute infection. The amount of TZ collagen correlates with the frequency of TGF-β+ macrophages which, in turn, correlates with the frequency of HIV RNA-producing cells. We conclude that immediate initiation of ART may have limited impact in preventing or reversing collagen accumulation in LNs.

## Introduction

HIV replication occurs primarily in lymphatic tissues (LTs) and is associated with a significant inflammatory response that includes deposition of collagen in the parafollicular T cell zone (TZ)^1-9^. As a consequence, the fibroblastic reticular cell network (FRCn) is disrupted, impairing access of naïve T cells to the homeostatic cytokine interleukin 7 (IL-7) produced by FRCn and leading to significant and sustained loss of T cells^8,10^. In a non-human primate model of SIV infection, increased TZ collagen deposition and FRCn damage was mediated by transforming growth factor β (TGF-β)+ T regulatory cells and occurred in the first weeks of infection following sexual mucosal transmission^11^.

We undertook the study reported here to determine if initiation of antiretroviral therapy (ART) in the earliest stages of acute HIV infection might block viral replication and associated immune activation to reduce TZ fibrosis. We studied participants in the RV254 cohort in Bangkok, Thailand, where individuals considered at high risk for HIV infection were screened for the presence of HIV RNA and ART was initiated as soon as it was detected. We studied LTs from before and during ART in this cohort to determine the rate of collagen deposition, the process by which it is formed, and if very early ART inhibited or reversed the process.

## Results

### Cohort description

We sampled peripheral blood and obtained an inguinal lymph node (LN) from 71 participants (2 females and 69 males) enrolled in the Thai Red Cross RV254 study^12^. Individuals were divided into two groups: the first group composed of 40 individuals who were immediately biopsied at the time of diagnosis of acute HIV infection a median of 5 days (quartiles 4, 7) after diagnosis. ART was initiated immediately after biopsy. The second group was composed of 31 individuals who deferred biopsy until after a median of 49.7 weeks (quartiles 47.9, 120.9) of ART. The specific clade of infecting virus was determined for 45 of the participants and, of those, 39 were HIV-1 CRF_A/E, 2 were non-typeable, and 4 were HIV-1 CRF A/E/B recombinants. Of the 40 that were immediately biopsied at the time of diagnosis, 3 individuals (1 each diagnosed in Fiebig 3, 4, and 5) were biopsied a second time between 46 and 48 weeks of ART. Of the 31 individuals that deferred biopsy, 2 individuals had 2 biopsies; both were identified in Fiebig 3. For one of the participants, the first sample was obtained 26 weeks after starting ART and the second sample was obtained at 97 weeks. For the other individual, the first sample was obtained 143 weeks after the start of ART and the second after 242 weeks.

Participants in both groups were stratified into five groups based on Fiebig stage (1-5) of HIV infection at time of diagnosis^13^ (**Supplementary Table 1)**. As expected, there was a rapid exponential increase in plasma HIV RNA, depending on Fiebig stage at the time of diagnosis, from a median plasma viral load (quartiles) of 15,140 (9,601, 54,325.25) HIV RNA copies/ml in Fiebig stage 1 to a peak median viral load in Fiebig stage 3 of 4,247,062 (1,333,265, 8,865,340) HIV RNA copies/ml. The median (quartiles) CD4+ T cell count at the time of diagnosis was 570 (455.5, 609.25) cells/µl in Fiebig stage 1, 293 (158, 339) cells/µl in Fiebig stage 2, 345 (244, 452.5; *n*=31) cells/µl in Fiebig stage 3, 424.5 (246, 460.25; *n*=8) cells/µl in Fiebig stage 4, and 298 (266, 374; *n*=5) cells/µl in Fiebig stage 5 (**Supplementary Table 1)**. For comparison, we recruited 13 individuals without HIV, who were part of the RV254 study, being screened regularly for HIV infection and obtained a LN biopsy under the companion RV304 protocol.

### Inflammatory damage to LNs in acute infection

We have previously shown that the TZ of LNs in HIV-infected people have varying degrees of inflammatory damage and accumulation of collagen fibers that is indicative of a fibrotic process occurring^5,7,8,10,14-19^. We measured the amount of collagen in the TZ by staining tissues with antibodies directed against collagen 1 and used quantitative image analysis (QIA) to isolate the TZ and calculate the percent area with collagen as a measure of fibrotic changes to the TZ, as we have done in the past^5,8,9,16^. Among the HIV-negative participants, the median (quartiles) percent TZ area with collagen was 15.9% (13.1%, 19.4%). However, in Fiebig 1 (**Fig. 1 A, 1B**) the median (quartiles) percent area of the TZ with collagen for those biopsied at time of diagnosis (group 1) was 26.9% (24.5%,29.6%), 25.1% (22.7%, 27.0%) in Fiebig 2, 24% (20.8%, 27.1%) in Fiebig 3, and 22.3% (21.2%, 27.2%) in Fiebig 4/5 (**Fig. 1C**). The difference between HIV-negative and Fiebig 1 was significant (*p*=0.0077, multivariable linear regression, **Supplementary Table 2**) and there was no difference in TZ collagen between Fiebig stages at time of diagnosis (**Supplementary Table 3**). We had also hypothesized that initiation of ART during acute infection would limit TZ collagen formation. However, we found that among the individuals who initiated ART at the time of acute infection and who deferred biopsy until after a median of 49 weeks of ART (group 2), there was no difference in TZ collagen when compared to measures of collagen in group 1 who were sampled at the time of diagnosis and before ART(**Fig. 1D, Supplementary Table 3**). Next, we compared the area of the TZ with collagen to duration of ART, after adjusting for Fiebig stage at enrollment and age at biopsy, to determine if there was any impact of ART on collagen formation and we found no relationship (*p*=0.740, **Fig. 1E**). Collectively, these data show that fibrotic changes in LN architecture occurs immediately after virus acquisition, is persistent, and does not resolve with early ART.

**Figure 1.**
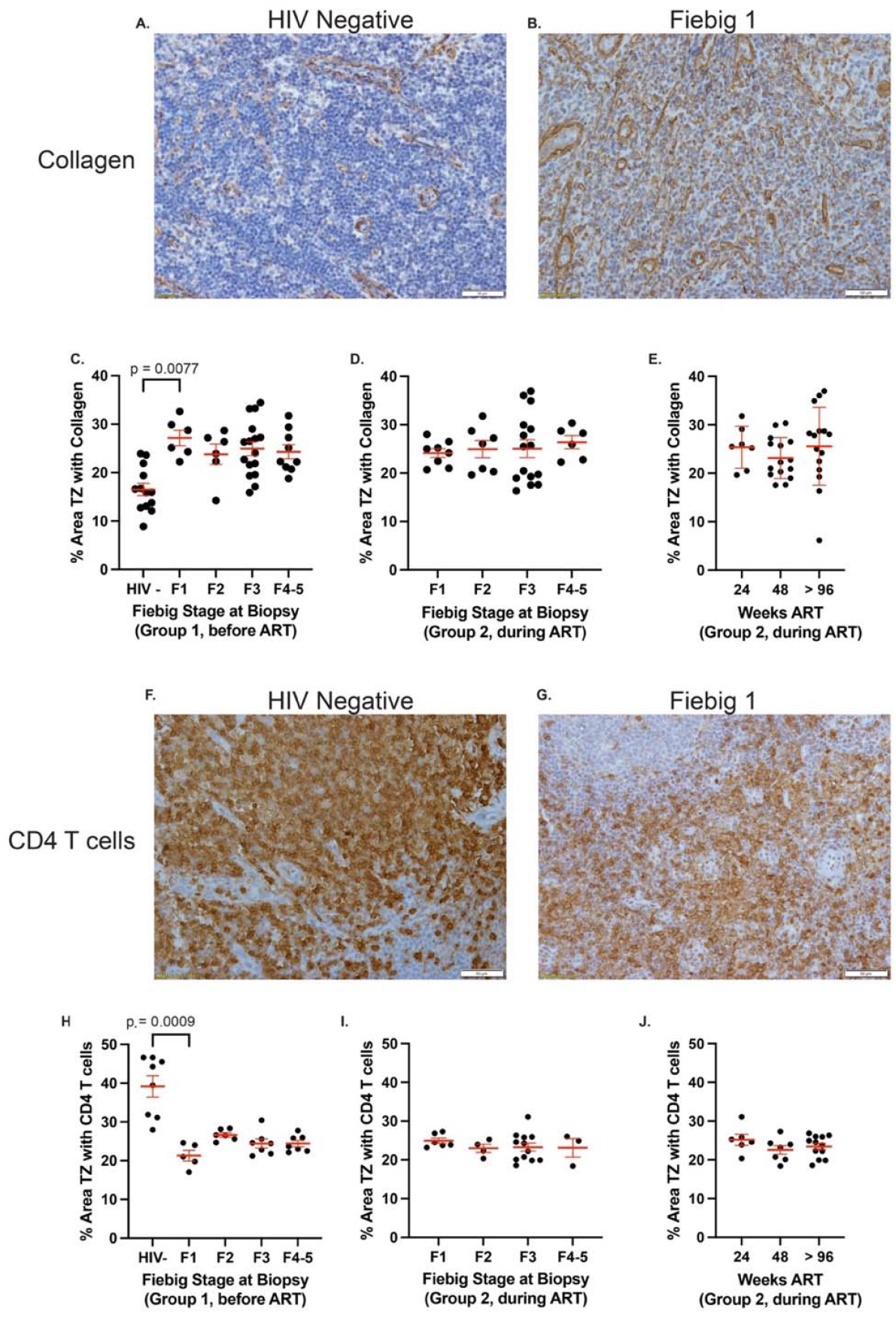
Significant TZ collagen accumulation occurs by Fiebig 1. In A, we show a representative section stained with antibodies against collagen 1 from a person without HIV in RV254. In B, we show a participant in Fiebig 1 demonstrating a significant increase in TZ collagen. In C, we show the quantitative image analysis of the percent area of the TZ that stains positive for collagen in the participants sampled during acute infection. There is a significant increase in TZ collagen in the participants sampled during Fiebig 1 compared to the people without HIV (*p*=0.0077, multivariable linear regression). In D and E, we show that there is no change in collagen after a mean of 49.7 weeks of ART. In panels F-J, we show the same type of analysis for LN CD4+ T cells demonstrating the rapid loss of CD4+ T cells in the LN and the fact that they are not restored with ART. There is a significant decrease in CD4+ T cells in the participants sampled during Fiebig 1 compared to individuals without HIV (*p*=0.0009, multivariable linear regression).

We next measured the impact of collagen deposition in the FRCn on the LN CD4+ T cell population using antibodies directed against CD4 and QIA to determine the frequency of TZ CD4+ T cells using previously developed and validated methods that quantified the percent area of the TZ with cells staining positive for CD4^5,8^. QIA revealed a significant loss of TZ CD4+ T cells as early as Fiebig 1 (**Fig. 1F, 1G**). In individuals without HIV, CD4+ T cells occupied a median (quartiles) of 41.9% (31.7, 45.8) of the TZ whereas in those biopsied at time of diagnosis in Fiebig 1 it was 21.0% (19.8%, 24.0%), 26.5% (25.8%, 27.8%) in Fiebig 2, 24.4% (22.3%, 25.1%) in Fiebig 3, and 23.7% (22.8%, 25.9%) in Fiebig 4/5 (**Fig. 1H**). The difference between HIV-negative and Fiebig 1 was significant (*p*=0.0009, multivariable linear regression, **Supplementary Table 2**). At the time of diagnosis, the area of the TZ with CD4+ T cells for individuals in Fiebig 1 was also significantly lower than compared to individuals in Fiebig 2 (adjusted *p*=0.005, multivariable linear regression; **Supplementary Table 4**). For individuals sampled after prolonged ART (group 2), there was no significant difference in the area of the TZ with CD4+ T cells dependent on Fiebig stage at ART initiation (**Supplementary Table 4**). Similarly, area of the TZ with CD4+ T cells was not associated with duration of ART (*p*=0.642) (**Fig. 1I, 1J**). Thus, there was substantial LN T cell depletion in the earliest stages of HIV infection and ART had no effect on restoring that population over time.

### Significant induction of TGF-β in acute HIV infection

Based on studies with a non-human primate model of SIV infection in which TGF-β was shown to play an important role in TZ collagen deposition^11^, we wanted to study this further in HIV infection. We first measured plasma levels of TGF-β in four different groups; in the HIV-Thai individuals, in group 1, sampled during acute infected and before ART was begun, in group 2 individuals who were started on ART during acute infection and sampled after 24-120 weeks of ART, and people with chronic HIV infection (**Fig. 2A**). These data show a significant increase in plasma TGF-β during acute infection that is persistent regardless of length of ART. We next used IHC with antibodies against TGF-β to determine the frequency of LN TGF-β+ cells and plot these frequencies against duration of ART (**Fig. 2B**) to demonstrate persistent levels of cells expressing TGF-β in LNs. Finally, the frequency of HIV RNA+ cells (vRNA+) by RNAscope in LNs was determined at diagnosis (group 1) and during ART (group 2) demonstrating persistent virus production in most individuals on ART (**Fig. 2C**). We had sufficient tissues to measure (vRNA+ cells in 15 of the group 1 individuals (i.e., sampled in acute infection prior to ART), and we found a significant correlation between vRNA+ cells and TGF-β+ cells. As the frequency of vRNA+ cells increased so did the frequency of cells expressing TGF-β (**Fig. 2D**, *p*=0.0077, multivariable linear regression, **Supplementary Table 5**).

**Figure 2.**
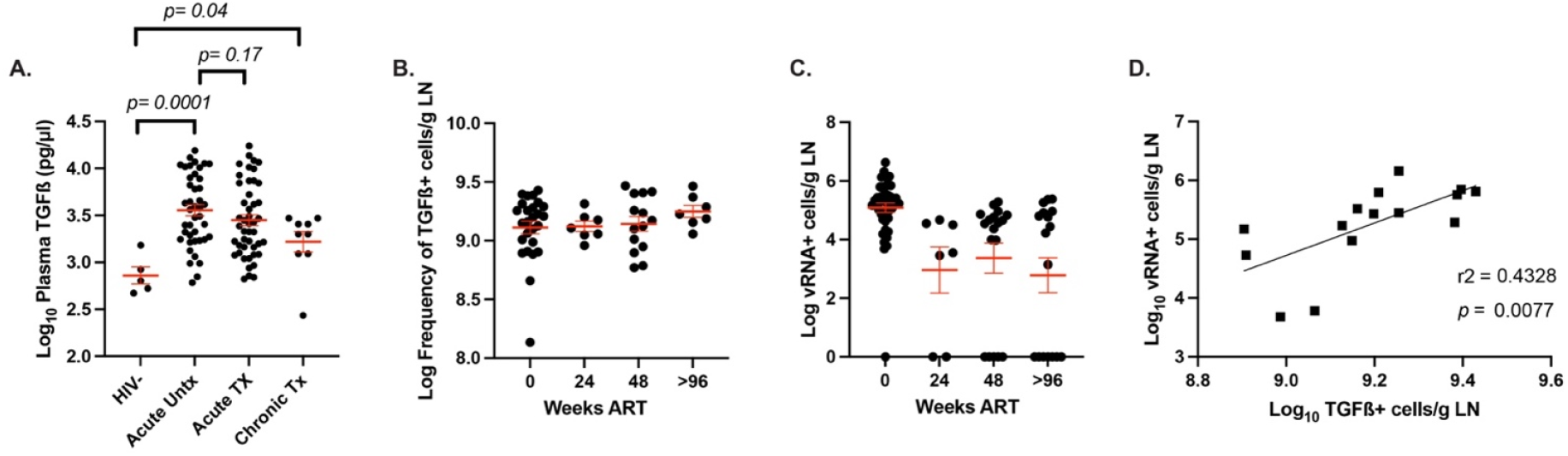
TGF-β expression in plasma and LNs. A. plasma levels of TGF-β are significantly increased in HIV infection and remain elevated even with ART. B. The frequency of cells expressing TGF-β in LN was measured before and during ART showing a pattern of persistent TGF-β production in LNs despite ART. C. The frequency of vRNA+ cells in LNs was determined at diagnosis (group 1) and during ART (group 2) demonstrating persistent virus production in most individuals on ART. D. There is a significant relationship between the frequency of TGF-β+ and HIV vRNA+ cells in LN. Statistics from multivariable linear regression, results from model adjusted for age at biopsy given in **Supplementary Table 5**.

Given the association between virus production and TGF-β in LNs we next sought to determine the cellular sources of TGF-β. In SIV-infected macaques, T regulatory cells were identified as a primary source of TGF-β, and we were interested to know if the same was true in HIV infection. We used double-label IHC to evaluate multiple cell types that have been associated with TGF-β production (**Supplementary Tables 6 and 7**) and found that most cells producing TGF-β were macrophages and that, in contrast to what was seen in acute SIV infection^16^, only ∼10% of TGF-β+ cells were T regulatory cells. In addition, there were small populations of endothelial cells and neutrophils that were TGF-β+. We stained LN sections from a Thai individual without HIV for comparison to LN sections from individuals in the acute and chronic stages of HIV infection using antibodies directed against TGF-β, CD31 to mark endothelial cells, and CD68 to mark macrophages (representative images shown in **Fig. 3)** In the HIV-negative individual, most TGF-β+ cells were endothelial cells lining vessels and relatively few macrophages were TGF-β+. (**Fig. 3A**). In contrast, in Fiebig 1 there was an expansion of cells producing TGF-β, with most coming from macrophages (**Fig. 3B**). In Fiebig 3 and chronic infection, TGF-β production was decreased compared to Fiebig 1 but was still predominantly from macrophages (**Fig. 3 C, 3D**).

**Figure 3.**
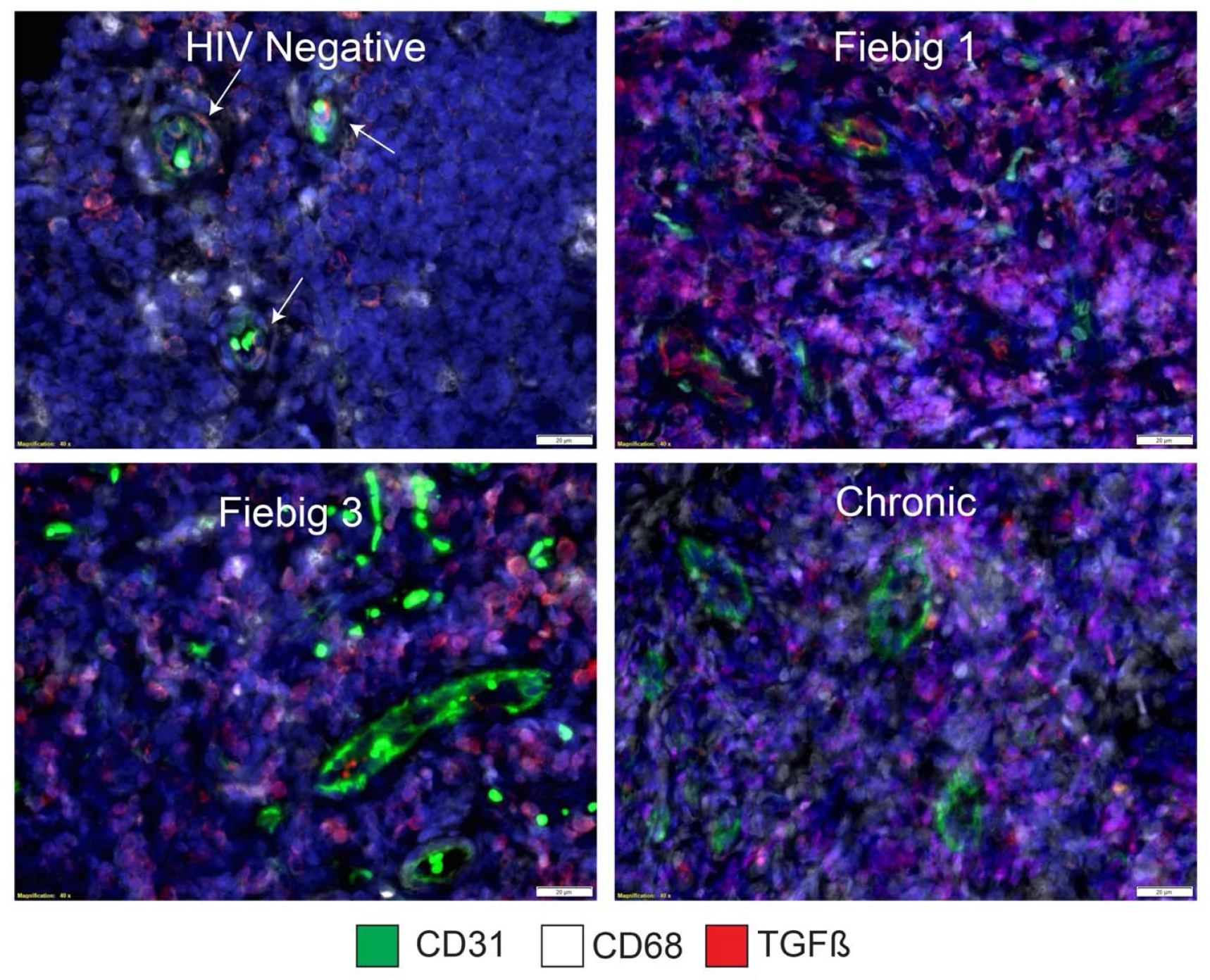
TGF-β in lymph nodes in HIV infection is associated with production in macrophages. In a Thai individual without HIV, a section of LN is stained with antibodies against TGF-β (red), CD68 to mark macrophages (white), and CD31 to mark epithelial cells (green) (upper left panel). TGF-β+ cells are relatively infrequent and are either epithelial cells (white arrows) or CD68-cells. In Fiebig 1 (upper right panel) there is a significant increase in TGF-β cells and virtually all of them are also CD68+ indicating production from macrophages (appearing pink in the figure). This pattern continues into Fiebig 3 and chronic HIV infection.

We were interested to know the phenotype of macrophages producing TGF-β in HIV infection and stained tissues with antibodies directed against TGF-β, CD68, and CD163 to distinguish the more proinflammatory macrophage (CD68+/CD163-) from the more anti-inflammatory macrophage (CD68+/CD163+). In **Fig. 4A**, we show a representative image from a LN from Fiebig 1 where virtually all of the TGF-β+ cells are macrophages of the proinflammatory phenotype (CD68+/CD163-). However, by Fiebig 3 (approximately 2 weeks later) we see the opposite pattern where most of the TGF-β+ cells are CD68+/CD163+ (**Fig. 4B**), which persists into chronic infection (**Fig. 4C**).

**Figure 4.**
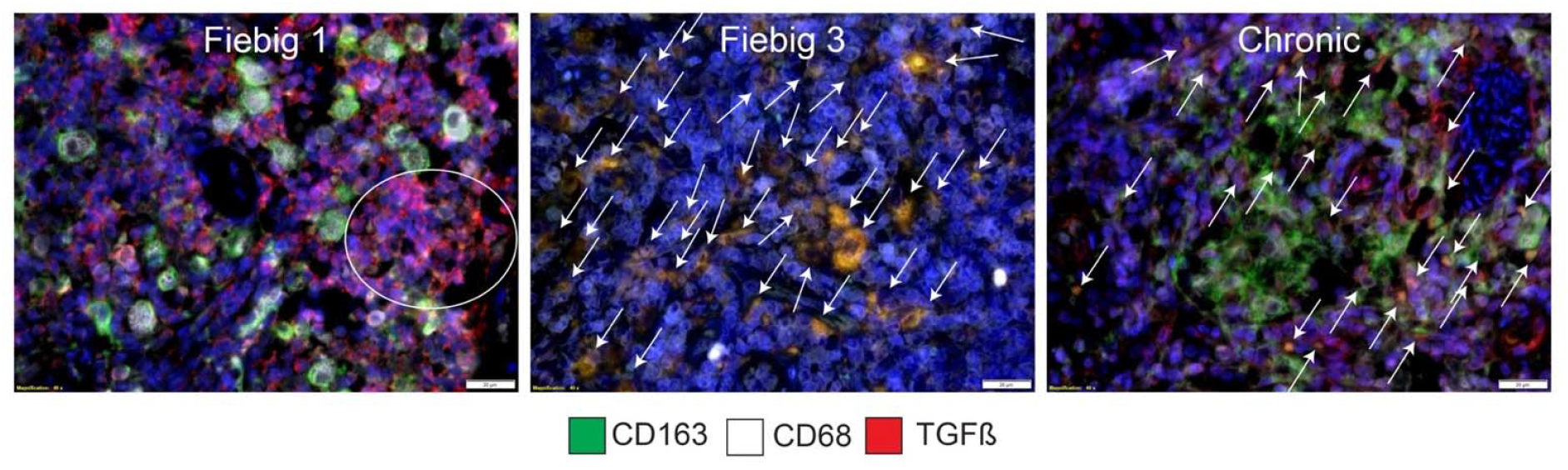
Phenotype of macrophage producing TGF-β. We stained LN sections from individuals sampled in Fiebig 1, Fiebig 3, and treated chronic infection with antibodies against CD68 (white) to identify macrophages, CD163 (green) to distinguish pro-inflammatory macrophages (CD68+/CD163-) from anti-inflammatory macrophages (CD68+/CD163+), and TGF-β (red). In Fiebig 1 (left panel), the majority of macrophages producing TGF-β are of the proinflammatory phenotype staining CD68+/CD169-. The white circle indicates an area with a high frequency of TGF+/CD68+/CD163-cells that are pink in color. The anti-inflammatory macrophages (green/white) are not TGF-β+. In Fiebig 3 (middle panel) virtually all of the TGF-β cells are of the anti-inflammatory phenotype and appear orange in color. White arrows point to the TGF-β+/CD68+/CD163+ cells. In treated chronic HIV infection, the majority of TGF-β cells are of the anti-inflammatory phenotype (white arrows).

## Discussion

Our original hypothesis was that ART started in the earliest stages of HIV infection would prevent or reverse collagen accumulation and preserve CD4+ T cell populations. Surprisingly, this was not the case. We show that there was significant accumulation of TZ collagen even in the earliest stages of HIV infection and that initiation of ART in the earliest detectable stage of HIV infection (Fiebig 1) had no impact on limiting or reversing the amount of collagen in LNs or restoring LN populations of CD4+ T cells.\

We further show that macrophages play a key role in this abnormal collagen accumulation in this earliest stage of HIV infection through production of TGF-β that was correlated with the frequency of vRNA+ cells, consistent with virus production as the stimulus for macrophage production of TGF-β. We have previously shown that HIV producing resting CD4+ T cells dominate virus production in the TZ in Fiebig 1 and that these virus-producing cells persist through two to five years of ART^20^.

We thus conclude that LN virus production provides the stimulus for macrophages to produce TGF-β both before and after initiating ART, pointing to the potential beneficial impact of interventions that target TGF-β and limit virus production that would also limit the pro-fibrotic destruction of LN structure that support CD4+ T cell populations^5,7-10,14,15,17,18,21^.

## Methods

### Sex as a biological variable

Sex was considered a biological variable. Our protocol was written to include any eligible participant regardless of sex or gender, and our recruitment efforts followed that principle. The participant population of this study generally reflected the demographics of PWH in Minnesota. Most study participants were male.

### Plasma HIV RNA

Plasma HIV RNA was measured using either the Roche Amplicor HIV-1 Monitor Test v1.5 or the Roche COBAS AmpliPrep/COBAS TaqMan HIV-1 Test v2.0 (Roche Diagnostics, Branchburg, New Jersey, USA). Lower limits of detection were 50 and 20 copies/ml, respectively.

### CD4+ T cell counts

CD4+ T cell counts were measured by either single- or dual-platform flow cytometry (Becton-Dickinson Biosciences, San Jose, California, USA).

### Tissue fixation and sectioning

Tissues were fixed in fresh 4% paraformaldehyde (Electron Microscopy Sciences) in PBS for 24 hours at room temperature. Tissues were washed in 80% ethanol for 5 minutes, 3 times before storing at 4°C until embedding. Five-micron sections were cut and mounted on EPIC Plus slides (Epic-Scientific).

### In situ hybridization

Slides were dewaxed at 60°C for 1 hour before transferring to xylenes for 10 minutes (twice), and 100% ethanol 5 minutes (twice). RNAscope was performed as done previously^22^ using the RNAscope 2.5HD Reagent Kit-RED (Advanced Cell Diagnostics). Briefly, slides were boiled for 10 minutes in Target Retrieval Reagent, cooled in DI water and dipped 10 times in 100% ethanol. Tissues were circled with an ImmEdge hydrophobic pen (Vector Laboratories), before adding Protease Plus, (diluted 1:10 in cold PBS) for 20 minutes in the 40°C humidified HybEZ oven. Slides were then washed in DI water before adding the HIV RNA target probe (vRNA anti-sense probe, ACD catalog number 446551) for 2 hours. Slides were rinsed twice with 0.5X wash buffer before adding AMP 1 for 30 minutes at 40°C. Following 2 more washes, AMP 2 was added for 15 minutes at 40°C. Slides were washed twice before AMP 3 was added for 30 minutes at 40°C. Following 2 more washes, AMP 4 was added for 15 minutes at 40°C. Slides were washed twice before adding AMP 4 for 30 minutes at room temperature. Again, following 2 more washes, AMP 6 was added for 15 minutes at room temperature. Slides were washed twice. Fast Red Chromogen (Biocare Medical) was added 1:1500 for 2 minutes, rinsed in TBST, before counterstaining in CAT hematoxylin (Biocare Medical), and blued in TBST for 1 minute.

### Immunohistochemistry

Slides were dewaxed, hydrated to water before antigen retrieval was performed with their appropriate buffer for 125°C for 30 seconds using a Declocking chamber (Biocare Medical). After cooling, tissues were circled with a hydrophobic pen and blocked with 5 minutes in 3% Hydrogen Peroxide, rinsed in TBST and blocked in Background Sniper (Biocare Medical) for 30 minutes. Primary antibodies were diluted in Divinci Green (Biocare Medical) and incubated overnight at 4°C. Tissues were rinsed 3 times in TBST before adding their secondary HRP antibodies per manufacturers directions (Origene). Slides were developed with Immpact DAB (Vector) and counterstained in hematoxylin. Slides were dehydrated, followed by 5 minutes in xylenes (twice) and mounted in permount (Fisher Chemical).

### Multiplex immunofluorescence

Slides were prepared as above. Antigen retrieval was done in DIVA buffer (Biocare Medical), followed by a 10-minute incubation step in pK solution in a 40°C waterbath. Slides were rinsed in TBST 3x for 5 minutes before blocking in Sniper blocking solution for 30 minutes. TGF-β1 (Biotechne) 1:300 diluted in Opal Diluent (Akoya biosciences) overnight at 4°C. After rinsing in TBST 3X, anti-goat HRP (Origene) was added according to manufacturer’s instructions. Opal 570 (Akoya) was diluted 1:300 and added to slides for 10 minutes in the dark. Slides were stripped in their next antibody’s appropriate buffer for 10 minutes in a microwave at 20% power. After cooling, staining was repeated with the next antibody, anti-mouse/anti-rabbit HRP (Akoya), and OPAL 488 or OPAL 647 (Akoya). DAPI was added for 5 minutes before mounting in Aqua-Poly/Mount (Polysciences).

### Image acquisition

IHC- and ISH-stained slides were imaged using brightfield whole-tissue scanning by an Aperio Versa 8 (Leica Biosystems) with a 20x/0.80NA objective. Digitized images (Leica.SCN File Format) were viewed and annotated using Aperio eSlide Manager. Monochrome images for IFA-stained slides were captured for each fluorophore using an Olympus DP80. A pseudo color was applied using the CI Deconvolution algorithm (Olympus) for each marker and fused into a composite image for viewing.

### Quantitative image analysis

Image processing and analysis for Collagen I stained slides was performed using the Area Quantification module of Halo Image Analysis Platform version 3.4.2986 (Indica Labs, Inc). Using the Area Quantification module, the percentage of tissue area positive for Immpact 3,3-diaminobenzidine (Vector Laboratories) staining was generated using the optical density for DAB chromogen and the Hematoxylin counterstain as a proportion of the tissue area analyzed (HALO® Image Analysis Platform, Computer Software 3.4.2986, Albuquerque, NM: Indica Labs Inc. 2021). For CD4 area quantification analysis, Photoshop (Adobe) was utilized to mask empty space and artifacts in the tissue white. Tissue stained positive with DAB chromogen was set to black by intensity thresholding and then rendered as a binary image after being exported to ImageJ. To find the percent area of positive staining the binary image was then analyzed using the Measure tool and results are expressed as percent area of positive staining per total region of interest. To quantify TGF-β positive cells, a binary image was rendered in ImageJ as described above and a value found using the Analyze Particles tool. This value was then used to calculate the positive cells per gram using the area in um^2^ measured by Aperio eSlide Manager^5,7,8,23^.

### Statistical analysis

Continuous values were summarized using medians and quartiles. Categorical variables were summarized using proportions. Area of the TZ with collagen and CD4+ T cells were arcsine transformed prior to statistical analysis. Frequency of vRNA+ and TGF-β+ cells per gram were log base 10 transformed. Due to small sample sizes, Fiebig stages 4 and 5 were grouped together for analyses.

To account for correlation between measurements taken from the same individual, generalized estimating equations (GEE) were used to obtain parameter estimates and robust standard errors using independence working covariance matrices. Outcome variables in these models were area of TZ with collagen and area of TZ with CD4+ T cells while independent variables included the interaction between Fiebig stage at enrollment and measurement timepoint (prior to ART, during ART), age at biopsy, and number of weeks on ART. For the analyses of the association between area of the TZ with collagen and CD4+ T cells and number of weeks on ART only age at biopsy, Fiebig stage at enrollment, and number of weeks on ART were included in the models. A linear regression model was used to assess the association between vRNA+ and TGF-β+ cells per gram, controlling for age at biopsy. Linear regression models were used to compare area of the TZ with CD4 cells and collagen between HIV-negative individuals and individuals starting ART in Fiebig stage 1, while adjusting for age at biopsy. Unadjusted p-values and Holm adjusted p-values are presented for each analysis, as appropriate.

### IRB approval

All samples analyzed in this study were obtained with the written consent of participants using protocols approved by Institutional Review Boards/Ethical Committees at the Walter Reed Army Institute of Research (approval number 1494), Chulalongkorn University, Bangkok, Thailand (approval number 220/51) and the University of Minnesota (approval number 1604M87147). The archived tissues used as historical chronic controls were obtained from participants in an IRB-approved study at the University of Minnesota (approval number 00009301, ClinicalTrials.gov (NCT04311177).

## Data availability

Deidentified data are available from the corresponding author upon reasonable request.

## Author contributions

All authors made substantial contributions to this work. TWS, DCD, SV, and JA conceived the study. CPS, SS, NP, and AS recruited and managed participants. JGC assisted with training for lymph node biopsy in a research setting. TWS, DCD, and JA designed experiments. JA, CD, and GW completed the tissue analyses. KE, TWS, SV, and EH wrote the paper. EH conducted the statistical analyses.

## Acknowledgments

Antiretroviral therapy for RV254/SEARCH 010 participants was supported by the Thai Government Pharmaceutical Organization, Gilead, Merck and ViiV Healthcare. Material has been reviewed by the Walter Reed Army Institute of Research. There is no objection to its presentation and/or publication. The views expressed are those of the authors and should not be construed to represent the positions of the U.S. Army or the DoD. The investigators have adhered to the policies for protection of human participants as prescribed in AR 70–25. We thank Jintanat Ananworanich for her contributions in the design and execution of this study.

