## Supplementary for "HIV lymphoid tissue fibrosis occurs in the earliest stages of acute HIV infection and is associated with macrophage-derived TGF-β"

| **Supplementary Table 1.** Demographic characteristics at the time of biopsy for the study cohort. Median (quartiles) are presented unless otherwise specified. | | | | | | | | | | | |
| --- | --- | --- | --- | --- | --- | --- | --- | --- | --- | --- | --- |
|  | Fiebig 1 | | Fiebig 2 | | Fiebig 3 | | Fiebig 4 | | Fiebig 5 | | HIV neg |
| Timing of biopsy | Immediate | Deferred | Immediate | Deferred | Immediate | Deferred | Immediate | Deferred | Immediate | Deferred | N/A |
| *n* | 6 | 8 | 6 | 7 | 19 | 12 | 5 | 3 | 4 | 1 | 13 |
| Age at first biopsy (yrs) | 23.5 (22.2, 24.8) | 27.5 (26.8, 32) | 25 (22.2, 28.5) | 30 (24, 32.5) | 25 (22, 31.5) | 30 (23, 36.5) | 26 (23, 34) | 24 (24, 26) | 32 (23.8, 39.2) | 29 | 31 (28, 33) |
| Male, *n* (%) | 6 (100%) | 7 (88%) | 6 (100%) | 7 (100%) | 19 (100%) | 11 (92%) | 5 (100%) | 3 (100%) | 4 (100%) | 1 (100%) | 10 (77%) |
| ART at first biopsy (wks) | 0.3 (0.1, 0.5) | 95.2 (50, 110.6) | 0 (0, 0.2) | 48 (35.9, 49.1) | 0.1 (0, 0.5) | 49.6 (43.6, 143.8) | 0.3 (0.3, 0.4) | 48.9 (36.8, 97.3) | 0.1 (0.1, 0.4) | 98.9 | N/A |
| CD4 at diagnosis (cells/µl) | 516 (392, 576) | 570 (456, 708) | 252 (171, 324) | 338 (178, 396) | 359 (249, 472) | 326 (188, 364) | 456 (438, 473) | 234 (208, 322) | 336 (282, 437) | 266 | N/A |
| CD4 at first biopsy (cells/µl) | 516 (392, 576) | 806 (653, 1004) | 252 (171, 324) | 609 (522, 706) | 359 (249, 472) | 568 (469, 710) | 456 (438, 473) | 516 (420, 554) | 336 (282, 437) | 341 (341, 341) | N/A |
| Log 10 plasma viral load at diagnosis (copies/ml) | 4.2 (4, 4.4) | 4.4 (3.9, 4.8) | 6 (5.5, 6.3) | 5.6 (5.4, 6.4) | 6.5 (6.1, 6.7) | 6.7 (6.1, 7.1) | 5.8 (5.2, 6.8) | 6.2 (5.9, 6.5) | 5.9 (5.7, 6.2) | 4.8 | N/A |
| Log 10 plasma viral load at first biopsy (copies/ml) | 4.2 (4, 4.4) | 1.3 (1.3, 1.3) | 6 (5.5, 6.3) | 1.3 (1.3, 1.3) | 6.5 (6.1, 6.7) | 1.3 (1.3, 1.3) | 5.8 (5.2, 6.8) | 1.3 (1.3, 1.3) | 5.9 (5.7, 6.2) | 1.3 | N/A |

**Supplementary Table 2.** Test of difference in estimated LT measures between biopsies collected from HIV-negative individuals and individuals starting ART in Fiebig stage 1. Estimates from separate multivariable regression models adjusted for age at biopsy are presented. Est Diff= Difference in measure for individuals in Fiebig stage 1 compared to HIV-negative individuals. CI=Confidence interval.

| **Test** | **Est. Diff.** | **95% CI** | **P-value** |
| --- | --- | --- | --- |
| Arcsine of area of TZ with CD4 cells | -0.26 | (-0.43, -0.08) | 0.0077 |
| Arcsine of area of TZ with collagen | 0.14 | (0.07, 0.22) | 0.0009 |

**Supplementary** **Table 3.** Test of difference in estimated arcsine transformed area of the TZ with collagen between groups defined by Fiebig stage at detection. Estimates from multivariable regression model adjusted for age at biopsy and duration of ART. Holm method for multiple comparisons used for p-value adjustment. Est=Estimated. Diff=Difference. CI=Confidence interval

|  |  |  |  | **P-values** | |
| --- | --- | --- | --- | --- | --- |
| **Comparison** | **Time** | **Est. Diff.** | **95% CI** | **Unadjusted** | **Adjusted** |
| Fiebig 1 – Fiebig 2 | Baseline | 0.042 | (-0.014, 0.098) | 0.1445 | 0.5780 |
| Fiebig 1 – Fiebig 3 | Baseline | 0.038 | (-0.003, 0.078) | 0.0711 | 0.4269 |
| Fiebig 1 – Fiebig 4/5 | Baseline | 0.039 | (-0.006, 0.084) | 0.0861 | 0.4306 |
| Fiebig 2 – Fiebig 3 | Baseline | -0.004 | (-0.057, 0.048) | 0.8686 | ≥0.9999 |
| Fiebig 2 – Fiebig 4/5 | Baseline | -0.003 | (-0.058, 0.053) | 0.9260 | ≥0.9999 |
| Fiebig 3 – Fiebig 4/5 | Baseline | 0.002 | (-0.038, 0.042) | 0.9294 | ≥0.9999 |
| Fiebig 1 – Fiebig 2 | On ART | -0.011 | (-0.056, 0.033) | 0.6218 | ≥0.9999 |
| Fiebig 1 – Fiebig 3 | On ART | -0.003 | (-0.051, 0.045) | 0.8965 | ≥0.9999 |
| Fiebig 1 – Fiebig 4/5 | On ART | -0.028 | (-0.065, 0.009) | 0.1376 | 0.8254 |
| Fiebig 2 – Fiebig 3 | On ART | 0.008 | (-0.043, 0.059) | 0.7582 | ≥0.9999 |
| Fiebig 2 – Fiebig 4/5 | On ART | -0.017 | (-0.061, 0.027) | 0.4533 | ≥0.9999 |
| Fiebig 3 – Fiebig 4/5 | On ART | -0.025 | (-0.072, 0.022) | 0.3026 | ≥0.9999 |

**Supplementary Table 4.** Test of difference in estimated arcsine transformed area of TZ with CD4+ T cells between groups defined by Fiebig stage at detection. Estimates from multivariable regression model adjusted for age at biopsy and duration of ART. Holm method for multiple comparisons used for p-value adjustment. Est=Estimated. Diff=Difference. CI=Confidence interval

|  |  |  |  | **P-values** | |
| --- | --- | --- | --- | --- | --- |
| **Comparison** | **Time** | **Est. Diff.** | **95% CI** | **Unadjusted** | **Adjusted** |
| Fiebig 1 – Fiebig 2 | Baseline | -0.065 | (-0.1, -0.03) | 0.0001 | 0.0005 |
| Fiebig 1 – Fiebig 3 | Baseline | -0.040 | (-0.08, 0) | 0.0350 | 0.1399 |
| Fiebig 1 – Fiebig 4/5 | Baseline | -0.043 | (-0.08, -0.01) | 0.0181 | 0.0903 |
| Fiebig 2 – Fiebig 3 | Baseline | 0.024 | (0, 0.05) | 0.0621 | 0.1863 |
| Fiebig 2 – Fiebig 4/5 | Baseline | 0.021 | (0, 0.04) | 0.0659 | 0.1863 |
| Fiebig 3 – Fiebig 4/5 | Baseline | -0.003 | (-0.03, 0.03) | 0.8342 | 0.8342 |
| Fiebig 1 – Fiebig 2 | On ART | 0.024 | (0, 0.05) | 0.0626 | 0.3757 |
| Fiebig 1 – Fiebig 3 | On ART | 0.018 | (-0.01, 0.04) | 0.1999 | 0.9997 |
| Fiebig 1 – Fiebig 4/5 | On ART | 0.022 | (-0.03, 0.07) | 0.3642 | 1.0000 |
| Fiebig 2 – Fiebig 3 | On ART | -0.006 | (-0.04, 0.02) | 0.6955 | 1.0000 |
| Fiebig 2 – Fiebig 4/5 | On ART | -0.002 | (-0.05, 0.05) | 0.9497 | 1.0000 |
| Fiebig 3 – Fiebig 4/5 | On ART | 0.005 | (-0.05, 0.06) | 0.8609 | 1.0000 |

**Supplementary Table 5.** Coefficient estimates for a linear model regressing log TGF-β+ cells/g LN on log vRNA+ cells/g LN. CI=Confidence interval.

| **Coefficient** | **Estimate** | **95% CI** | **p-value** |
| --- | --- | --- | --- |
| Intercept | -22.52 | (-43.90, -1.15) |  |
| Age at Bx | -0.01 | (-0.05, 0.03) | 0.6475 |
| log TGF-β+ cells/g LN | 3.05 | (0.65, 5.44) | 0.0169 |

**Supplementary Table 6: Antibodies used to determine phenotype of cells expressing TGF-β**

| Antibody | Company | Catalog # | Dilution | Antigen Retrieval buffer |
| --- | --- | --- | --- | --- |
| Collagen I | Abcam | 34710 | 1:200 | DIVA/ pk 4ul/ml |
| CD4 | Abcam | EPR6855 | 1:500 | DIVA |
| CD31 | DAKO | GA60 | 1:50 | Citrate |
| CD68 | Mouse | 0814 | 1:400 | Citrate |
| CD163 | Leica | NCL-L-CD163 | 1:25 | Citrate |
| TGF-β | Biotechne | AB-246-NA | 1:300 | DIVA/ pk 4ul/ml |

**Supplementary Table 7.** Antibodies used for IHC to detect the phenotype of cells producing TGF-β.

| **Antibody Name** | **company** | **dilution** | **Cell type** | **Frequency of TGF-β+ cell** |
| --- | --- | --- | --- | --- |
| CD44 | Abcam | 1:50 | Stromal cells | Negative |
| CD31 | DAKO | 1:50 | Endothelial cells | Few |
| CD11C | Novocastra | 1:20 | Dendritic cells | Negative |
| NK2GA | Abcam | 1:2000 | NK cells | Negative |
| CD15 | Biocare | 1:100 | Monocytes/ Granulocytes | Negative |
| EPCAM | Abcam | 1:100 | Epithelial cells | Negative |
| CD20 | Dako | 1:200 | B cells | Negative |
| Vimentin | Dako | 1:100 | Fibroblasts | Negative |
| CD68 | Dako | 1:400 | Macrophages | >75% |
| CD3 | Thermo | 1:300 | T cells | <5% |
| MPO | Thermo | 1:100 | Neutrophils | <5% |
| FOXP3/CD3 |  |  | T Regulatory cells | 10% |
